# Motor cortex is an input-driven dynamical system controlling dexterous movement

**DOI:** 10.1101/266320

**Authors:** Britton Sauerbrei, Jian-Zhong Guo, Matteo Mischiati, Wendy Guo, Mayank Kabra, Nakul Verma, Kristin Branson, Adam Hantman

**Affiliations:** Janelia Research Campus, Howard Hughes Medical Institute

## Abstract

Skillful control of movement is central to our ability to sense and manipulate the world. A large body of work in nonhuman primates has demonstrated that motor cortex provides flexible, time-varying activity patterns that control the arm during reaching and grasping. Previous studies have suggested that these patterns are generated by strong local recurrent dynamics operating autonomously from inputs during movement execution. An alternative possibility is that motor cortex requires coordination with upstream brain regions throughout the entire movement in order to yield these patterns. Here, we developed an experimental preparation in the mouse to directly test these possibilities using optogenetics and electrophysiology during a skilled reach-to-grab-to-eat task. To validate this preparation, we first established that a specific, time-varying pattern of motor cortical activity was required to produce coordinated movement. Next, in order to disentangle the contribution of local recurrent motor cortical dynamics from external input, we optogenetically held the recurrent contribution constant, then observed how motor cortical activity recovered following the end of this perturbation. Both the neural responses and hand trajectory varied from trial to trial, and this variability reflected variability in external inputs. To directly probe the role of these inputs, we used optogenetics to perturb activity in the thalamus. Thalamic perturbation at the start of the trial prevented movement initiation, and perturbation at any stage of the movement prevented progression of the hand to the target; this demonstrates that input is required throughout the movement. By comparing motor cortical activity with and without thalamic perturbation, we were able to estimate the effects of external inputs on motor cortical population activity. Thus, unlike pattern-generating circuits that are local and autonomous, such as those in the spinal cord that generate left-right alternation during locomotion, the pattern generator for reaching and grasping is distributed across multiple, strongly-interacting brain regions.

## Introduction

Reaching, grasping, and object manipulation play a central role in the lives of mammals with prehensile forelimbs. The musculoskeletal complexity of the limb poses a challenging control problem for the central nervous system, which must coordinate precisely-timed patterns of activity across many muscles to perform a wide diversity of tasks. The motor cortex is a brain region involved in the control of dexterous forelimb movement (*1–11*). In nonhuman primates, motor cortical lesions impair the coordination of the hand and fingers (*3*), and the activity of motor cortical neurons is closely linked to muscle activation, joint torques, and limb kinematics (*4, 8, 10*). In rodents, stimulation of motor cortex generates limb twitches (*12, 13*), chronic lesions impair dexterity (*14, 15*), and optogenetic inactivation blocks the initiation and execution of reaching (*16, 17*). However, several studies have concluded that motor cortex may play a fundamentally different role in rodents than in primates (*18*), such as tutoring other brain regions during learning (*19, 20*) or suppressing actions (*21*).

The function of motor cortex during reaching in nonhuman primates has been characterized by the dynamical systems model for reaching (rDSM) (*1, 2, 22*). This model proposes that the motor cortex and the arm are both dynamical systems that are linked through the firing rate output of motor cortex, r(t) (fig. 1a). Muscle activity, m(t) = G(r(t)), is determined by these firing rates, where G is a function describing how the lower motor centers transform the cortical output. Muscle activity, in turn, determines how the position and velocity of the joint centers in the arm, x(t), change over time. The arm kinematics evolve according to d/dt x(t) = F(m(t),x(t)), where F is a function describing the musculoskeletal mechanics. Motor cortical activity, r(t), evolves over time according to d/dt r(t) = h(r(t)) + u(t) as a result of two distinct influences: the local recurrent dynamics imposed by the architecture of motor cortex, which are described by a function h(r(t)) of the current state of motor cortex, and external input from other brain regions, described by a function of time, u(t). The external input, u(t), reflects the combined contribution of other brain regions, such as other cortical areas and thalamus, to changes in motor cortical firing rates. This input is not identical to the firing rates of the neurons in upstream brain regions; rather, it represents the effect those upstream firing rates have on firing rates in motor cortex.

**Figure 1:**
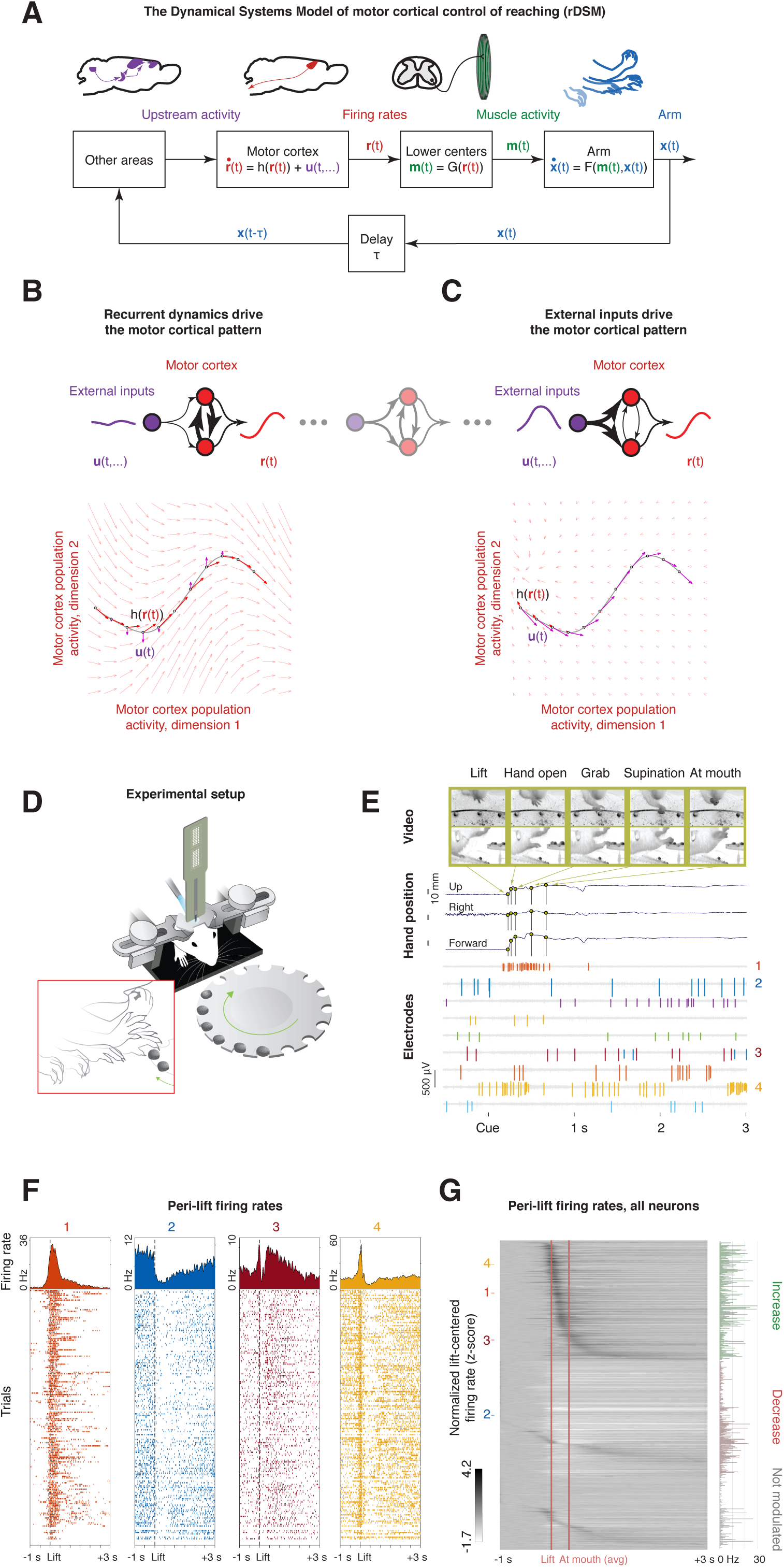
Motor cortex as a dynamical system controlling the arm. **a**, The dynamical systems model for motor cortical control of reaching (rDSM). In this model, the firing rates of motor cortical neurons, r(t), change as a result of two distinct influences. First, the local recurrent architecture intrinsic to motor cortex imposes a change, h(r(t)), based on the current firing rates. Second, brain regions outside motor cortex provide external input, u(t). Thus, the firing rates evolve according to d/dt r(t) = h(r(t)) + u(t). These firing rates control the muscle activation, m(t) = G(r(t)), through circuits in the lower motor centers, including the spinal cord. In turn, the muscles change the positions and velocities of the joint centers, x(t), through a function describing the musculoskeletal mechanics: d/dt x(t) = F(m(t),x(t)). Delayed sensory feedback from the arm ascends into the brain and influences the external inputs u(t), closing the loop. **b**, Generation of firing rate patterns if motor cortex were driven by strong recurrent dynamics. Under this possibility, changes in firing rate are dominated by recurrent dynamics, h(r(t)), arising from strong connections between neurons within motor cortex, while external inputs, u(t), have only a weak influence. The black line represents the neural trajectory, r(t). The light arrows in the background represent the vector field h(.). The bold red arrow represents this vector field evaluated at each point along the neural trajectory, r(t). The bold purple arrows represent the external input, u(t), evaluated at each time point along the neural trajectory, r(t). **c**, Generation of firing rate patterns if motor cortex were driven by strong external inputs. Under this possibility, changes in firing rate are dominated by strong external inputs, u(t), while the recurrent dynamics, h(r(t)), have only a weak influence. **d**, Experimental setup. Head-fixed mice reached for a pellet of food following an acoustic cue during recording and optogenetic perturbation of cortical activity. **e**, Raw video, electrophysiological recording, and mouse behavior on a single trial. Three-dimensional hand trajectories and the timing of each waypoint in the behavioral sequence were extracted from video using computer vision methods. **f**, Spike raster plots and peri-event time histograms for four example neurons, centered on lift. **g**, Average z-scored firing rates and mean firing rates for all motor cortical neurons (n = 19 mice, n = 39 sessions, n = 843 neurons). During prehension, most neurons exhibited increases (39%) or decreases (37%) in spike counts around lift (rank sum test with Benjamini-Hochberg correction, q < .05).

How do these two influences work together to generate the motor cortical output? One possibility is that the intrinsic circuitry of motor cortex contains all the machinery required to produce the time-varying output once it has been set to the appropriate initial condition; that is, the motor cortical dynamical system is largely autonomous from its inputs during movement execution, and the pattern results from the strong, local recurrent contribution, h(r(t)) (fig. 1b). Indeed, previous studies have focused on the role of recurrent dynamics in motor cortical pattern generation, and have suggested that external inputs may contribute by setting up the appropriate initial state during movement preparation (*23, 24*), while motor cortex is autonomous during movement execution (*2, 22, 25*). An alternative possibility, however, is that motor cortex is an input-driven dynamical system: that is, recurrent dynamics are not sufficient to generate the pattern, but that strong drive from external inputs, u(t), is required throughout the movement (fig. 1c). Here, we test these hypotheses by first validating a mouse system for motor cortical control of skilled reaching using electrophysiology and optogenetics, then using this experimental approach to show that the motor cortex is not autonomous during movement execution, but must be driven by strong external input.

### Cortical population recordings during reaching

While the rDSM is widely accepted as a model for cortical control of reaching in primates, it is unclear whether it is appropriate for rodents. Thus, we first tested whether the model describes the function of motor cortex in mice. One key requirement of the rDSM is that motor cortex must produce a time-varying output during reaching. In order to check this premise, we used a prehension task we developed for head-fixed mice (*16*), in which animals learned over several days to weeks to reach for and grab a food pellet at a memorized position and deliver it to the mouth following an auditory cue (fig. 1d). Using high-speed video and computer vision techniques, we captured the animals’ behavior and extracted the timing of waypoints indexing the stages of the movement (lift hand from the perch, open hand, grab food pellet, supinate hand, and bring hand to mouth), as well as three-dimensional hand position (fig. 1e; supplementary video 1). To study neural population activity during normal movement (supplementary fig. 1a), we used silicon probes to record spiking activity (mean 21 well-isolated units per recording) from a total of 843 neurons in forelimb motor cortex, which showed strong fluctuations in firing rate around the movement (fig. 1e-g). A majority of cells (646/843) were modulated before and during reaching, with a net rate increase for 330 and a decrease for 316 (fig. 1f; rank sum test on pre- and peri-lift spike counts with Benjamini-Hochberg correction, q < .05). While the responses of individual cells were highly consistent across trials (fig. 1f), we observed a wide diversity of patterns across neurons, including increases, decreases, and multi-phasic responses (fig. 1g). These activity patterns are qualitatively similar to those observed in the primary motor cortex of nonhuman primates performing dexterous behaviors (*8, 10*).

### Mouse motor cortex is a limb controller

Is this time-varying signal critical for producing the movement, as required by the rDSM? Two alternatives to the rDSM are that the cortical output must simply increase beyond a threshold to trigger movement, or that it must decrease below a threshold in order to ungate the movement. Neither of these threshold hypotheses is compatible with the rDSM: under the rDSM, a change in the motor cortical firing rate from the baseline to another constant value across a threshold will produce a change from one level of muscle activation to another constant value, but such a constant level of muscle activity cannot generate a complex, multi-step movement. In order to test the subthreshold hypothesis, we silenced excitatory neurons (and thus motor cortical output) by activating inhibitory neurons in VGAT-ChR2-EYFP mice (fig. 2a, left). This perturbation blocked the initiation of movement (fig. 2b, left, supplementary fig. 2a), ruling out the possibility that reaches are triggered when motor cortical output decreases below a threshold. In order to test the suprathreshold hypothesis, we activated intratelencephalic neurons (Tlx3-Cre X Ai32 mice; fig. 2a, center), which project to other motor cortical areas and the striatum, or pyramidal tract neurons, which project to the spinal cord and brainstem (Sim1-Cre X Ai32; fig. 2a, right). In both cases, the optogenetic perturbation blocked the initiation of reaching (fig. 2b, center and right; supplementary fig. 2b-c). These experiments rule out the possibility that motor cortical output to other motor cortical regions and the striatum, or to the spinal cord and brainstem, must simply increase above a threshold to trigger reaching. Taken together, these results suggest that reaching is not produced by shifts in the overall level of motor cortical output, but by a specific time-varying pattern.

**Figure 2:**
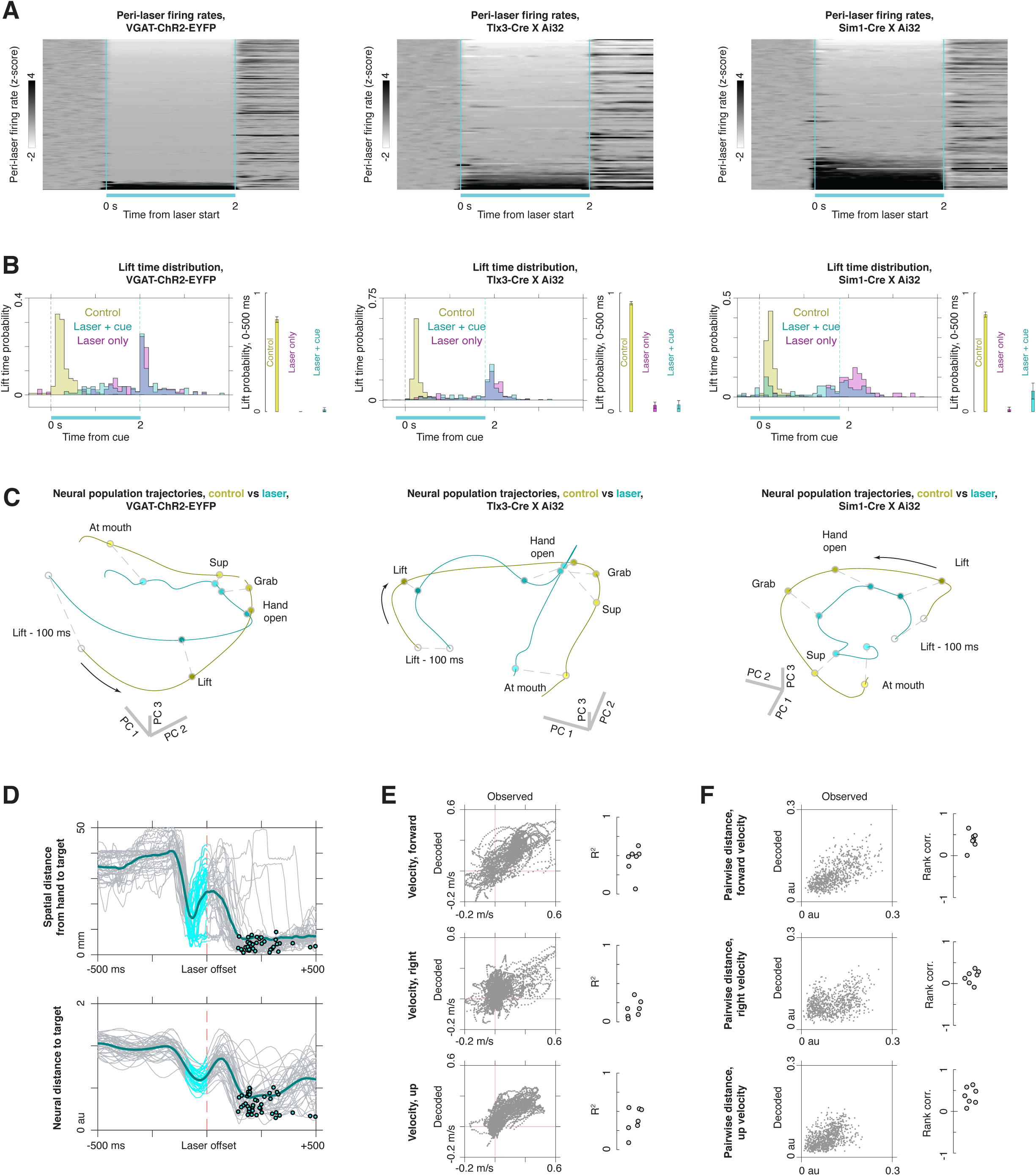
Motor cortex must generate a time-varying pattern throughout the movement. **a**, Firing rates of motor cortical neurons during optogenetic activation of inhibitory interneurons (left; VGAT-ChR2-EYFP mice), intratelencephalic neurons (center; Tlx3-Cre X Ai32 mice), and pyramidal tract neurons (right; Sim1-Cre X Ai32 mice). **b**, Distribution of lift times on control (yellow), laser + cue (blue), and laser-only (magenta) trials for VGAT-ChR2-EYFP mice (left), Tlx3-Cre X Ai32 mice (center), and Sim1-Cre X Ai32 mice (right). The histograms show data only for trials where a lift occurred. The insets to the right of the histograms show the probability that a lift occurred within the first 500 ms following the cue. **c**, Neural population activity from lift −100 ms to lift +350 ms on control (yellow) and laser (blue) trials, obtained using trial-averaged principal component analysis. Left, VGAT-ChR2-EYFP (n = 5 mice, n = 7 sessions, n = 144 neurons). Center, Tlx3-Cre X Ai32 (n = 3 mice, n = 7 sessions, n = 99 neurons). Right, Sim1-Cre X Ai32 (n = 3 mice, n = 5 sessions, n = 94 neurons). **d**, Upper: spatial distance from the hand to the target when the motor cortex is briefly inactivated in the middle of a reach in VGAT-ChR2-EYFP mice. The target is defined to be the average hand position at the time of grab on control trials. Lower: neural distance from the current state of cortex to the target state, defined to be the average value of the GPFA factors at grab time on control trials. Circles indicate grab times, and the blue regions indicate laser-on periods. **e**, Left: decoding of hand velocity from motor cortical spiking between 0 and 200 ms following the end of mid-reach optogenetic silencing. Each panel shows the decoded velocity in a single direction (forward, right, or up) for the dataset in **d**. Data are displayed only for test trials that were not used to train the decoder (see methods). Right: R2 values for decoded vs observed hand velocity for each dataset in which a midreach perturbation was applied (n = 4 mice, n = 7 sessions). **f**, Left: distance between observed and decoded velocity profiles for all pairs of trials in the dataset in **d**. Each point is one pair of trials. A positive correlation indicates that pairs of trials with similar observed velocity profiles also have similar decoded velocity profiles. Right: rank correlation between pairwise distance in observed and decoded velocity profiles for each dataset.

According to the rDSM, moving the arm to the target following a motor cortical perturbation would require motor cortex to regenerate the activity patterns that would normally drive reaching. Following the release of each motor cortical perturbation, we frequently observed kinematically normal post-laser reaches (fig. 2b; supplementary fig. 3a-b) (*16*). The reaches occurred with a shorter reaction time than on control trials following the VGAT and Tlx3 perturbations (supplementary fig. 2d-e), and they also occurred following laser stimulation in the absence of a cue (fig. 2b, magenta bars; supplementary fig. 2a-c). The neural activity patterns during reaches following each of the three perturbations largely recapitulated the patterns observed on control trials (fig. 2c, supplementary videos 2–3), and the neural state approached the average neural state at the time of grab from control trials (supplementary fig. 3d-h). The neural activity did not return to the initial state observed on control trials, but immediately generated the pattern for reaching following the end of the perturbation (supplementary fig. 4). In order to further compare the patterns of neural activity driving control and post-perturbation reaching, we trained a decoder to estimate the velocity of the hand from motor cortical firing rates. Even though the decoder was trained only on control trials (see methods), it was possible to decode the hand velocity from motor cortical activity following each of the three perturbations relatively well (supplementary fig. 5), further demonstrating that the neural patterns during post-laser reaches mostly recapitulated those occurring during normal reaching.

The rDSM maintains that the cortical output is coupled to the motor plant throughout the execution of movement, not merely at the initiation. When we briefly silenced motor cortex in the middle of a reach, the progression of the hand to the target halted, and following this inactivation, the motor cortical activity rapidly recovered, driving the hand to the target (fig. 2d). We were able to decode hand velocity from neural activity following the end of the perturbation (fig. 2e). Pairs of trials with similar post-perturbation hand observed velocities also had similar decoded velocities, suggesting that the output of the motor cortex can compensate for aberrant initial post-perturbation hand positions on a trial-by-trial basis (fig. 2f). Taken together, these results show that motor cortex is strongly coupled to the motor plant, and must generate a time-varying pattern throughout the entire movement sequence in order to control reaching, consistent with the rDSM.

### The motor cortical dynamical system is input-driven, not autonomous

After establishing that the rDSM (fig. 1a) describes the role of motor cortex during reaching in mice, we next sought to determine whether the motor cortical dynamical system is largely autonomous (fig. 1b) - that is, driven by its own local recurrent dynamics - or whether it is driven by strong external inputs (fig. 1c). In order to address this question, we observed that silencing motor cortex in VGAT-ChR2-EYFP mice fixes motor cortex to a constant state not only within the laser-on epoch on a given trial, but also across trials (supplementary fig. 1b; supplementary fig. 6a). This implies that the recurrent contribution to the dynamics will also be constant across trials during the laser-on epoch, and that trial-to-trial variability in neural activity following the release of the laser will reflect external inputs. Indeed, when we compared trials on which a post-laser reach occurred with trials with no reach (fig. 3a), we found that the two trial types started in the same initial state during laser-on, but rapidly diverged following the release of the laser (fig. 3b). The same was true at the single-trial level: on trials where the neural activity did not approach the average neural state at grab, the hand did not approach the pellet (fig. 3c). Because the initial neural state during the laser - and thus the contribution of recurrent dynamics to changes in firing rate - was the same for the reach and no-reach condition, the difference between conditions was driven by differences in external input (fig. 3d). The rDSM allowed us to directly estimate this difference in inputs by subtracting the firing rate derivatives of the two conditions (fig. 3e; see methods). The observation that motor cortex can diverge from the same initial state in different types of trials and drive different behaviors suggests that external inputs may play a key role in producing the activity pattern for reaching.

**Figure 3:**
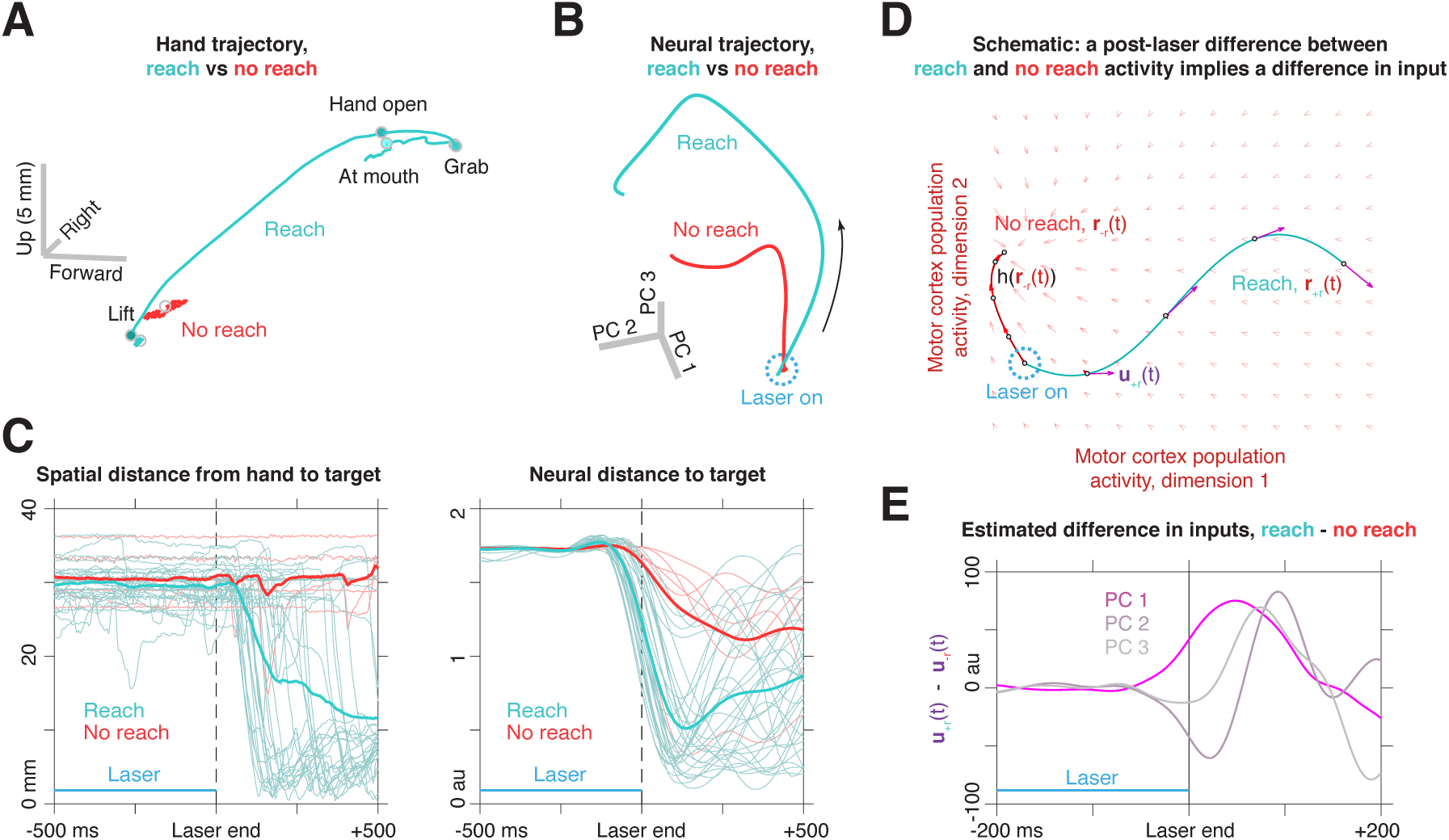
Distinct neural trajectories reflect the influence of different inputs. **a**, Hand trajectory for post-cortical-inactivation reaches in VGAT-ChR2-EYFP mice. Blue corresponds to trials in which a reach occurred within 700 ms following the end of the laser; time limits are −100ms to +425 ms from lift. Red corresponds to trials in which no lift occurred in the same window; because there is no lift, time limits are +100 ms to 625 ms from the end of the laser. **b**, Neural population activity aligned to the end of the laser for trials with (blue) and without (red) a post-laser reach. Time limits are 250 ms before the end of the laser through 250 ms after the end of the laser. **c**, Spatial (left) and neural (right) distance to target, centered on the end of the laser, for trials with (blue) and without (red) post-laser reaches. **d**, Schematic illustrating the conclusion that post-laser differences in neural trajectories implies a difference in external inputs. During the laser (dotted blue circle), the state of cortex has been set to the same value in trials with a reach, r_+r_(t), and trials without a reach, r_-r_(t). By the rDSM, the recurrent contribution to the dynamics, is also constant during the laser in the two conditions: r_+r_(t) = r_-r_(t). Thus, immediately following the end of the laser, differences in firing rate between the two conditions, r_+r_(t) and r_-r_(t), must be due to differences in external input, u_+r_(t) and u_-r_(t). **e**, Estimated difference in external inputs between reach and no reach conditions. The rDSM allows us to calculate the difference in inputs between the conditions, u_+r_(t) - u_-r_(t) (see methods). The difference for the first three dimensions of population activity (the first three principal component scores) are plotted.

If the motor cortex is strongly input-driven, then blocking or interrupting the input pattern should perturb both motor cortical activity and arm kinematics. In order to test this hypothesis directly, we implanted optical fibers above motor thalamus in VGAT-ChR2-EYFP mice. This enabled us to activate inhibitory terminals in this region, which has been shown to suppress activity in the projections to motor cortex (fig. 4a) (*26*). Thalamic inactivation at the start of the trial blocked the initiation of coordinated reaching (fig. 4b), and inactivation in the middle of the reach interrupted the progression of the hand to the target (fig. 4c, supplementary fig. 8). These results demonstrate that external inputs are required to control the hand throughout the entire movement.

**Figure 4:**
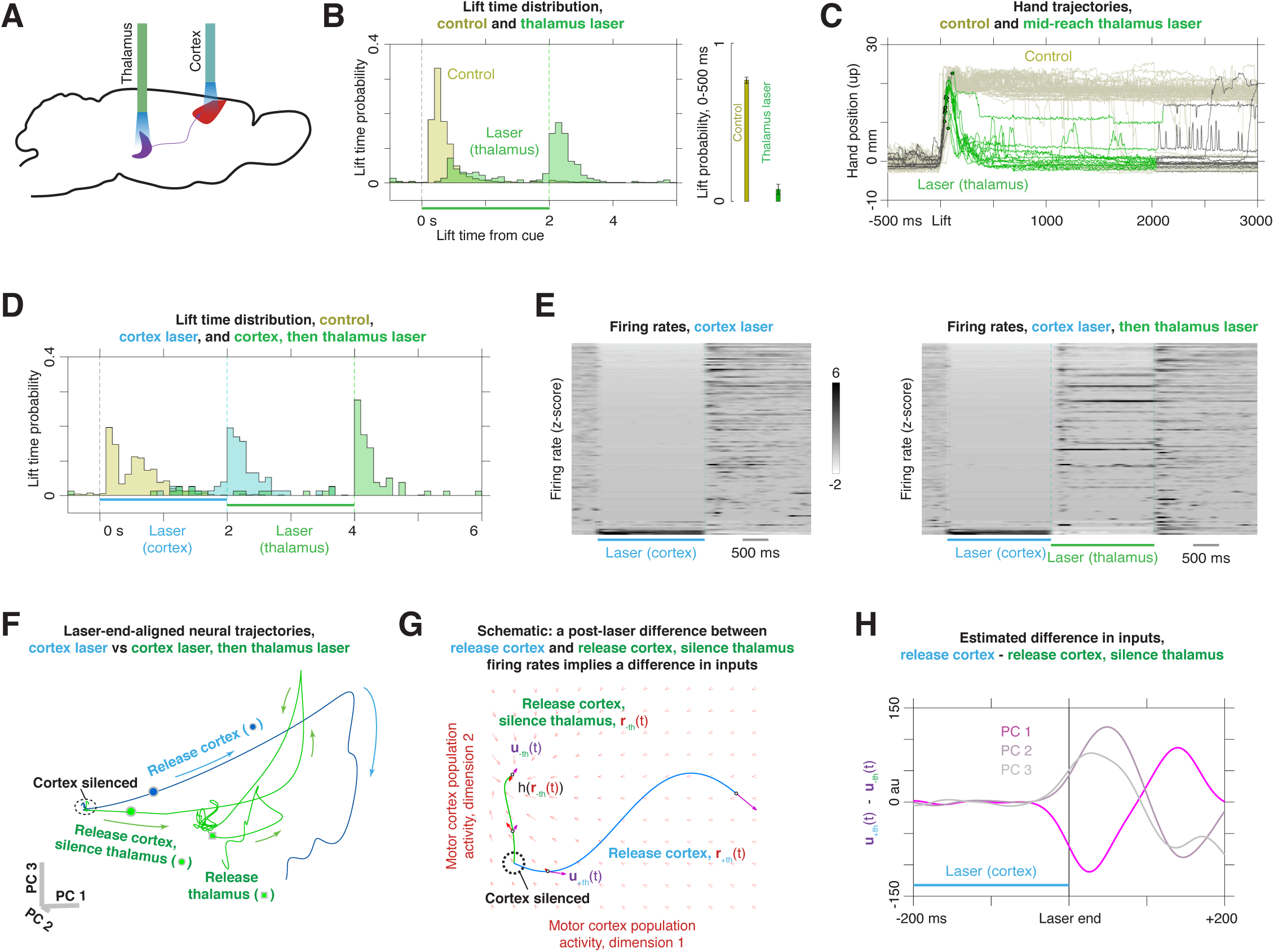
Strong external inputs drive the motor cortical pattern during reaching. **a**, Experimental schematic. One fiber was implanted in thalamus, and a second was positioned over motor cortex. **b**, Distribution of lift times on control trials (yellow) and on trials in which thalamus was inactivated for two seconds starting at the cue (green); n = 3 mice (VGAT-ChR2-EYFP), n = 12 sessions. Right inset shows the probability of a lift within the first 500 ms following the cue for cortex and thalamus inactivation trials. **c**, Hand position in the upper direction centered on lift on control trials (light yellow) and mid-reach thalamic inactivation trials (black; green indicates laser on) for a single dataset. Dots indicate the start of the laser. **d**, Lift times for control trials (yellow), cortical inactivation (blue), and sequential inactivation of cortex and thalamus (green); n = 3 mice (VGAT-ChR2-EYFP), n = 4 sessions. **e**, Firing rate Z-scores for all recorded neurons under inactivation of cortex alone (left) and sequential inactivation of cortex and thalamus (right); n = 3 mice, n = 4 sessions, n = 127 neurons. **f**, Population activity following the end of cortical inactivation for trials with cortical inactivation only (blue) and inactivation of thalamus after cortex (green). Plotting limits start 500 ms before the end of cortical inactivation and finish 500 ms after the cortical inactivation (blue trace) and 500 ms after the thalamic inactivation (green trace). Circles indicate the end of the cortical inactivation, and the square indicates the end of the thalamic inactivation. **g**, Schematic illustrating the effects of thalamic inactivation following cortical inactivation. During the cortical inactivation, firing rates on trials in which the thalamus will be inactivated, r_-th_(t), and trials on which it will not be inactivated, r_+th_(t), start in the same initial state (dotted black circle). By the rDSM, the recurrent contribution is the same in the two conditions: h(r_+th_(t)) = h(r_-th_(t)). Following the end of the cortical inactivation, the neural activity in the two conditions diverges due to a difference in external input: u_+th_(t) ≠ u_-th_(t). **h**, Estimated difference in external inputs between thalamus inactivated, u_-th_(t), and not inactivated, u_+th_(t), conditions. The rDSM allows us to estimate the difference in inputs between the conditions, u_+th_(t) - u_-th_(t) (see methods). The difference for the first three dimensions of population activity (the first three principal component scores) are plotted.

If external inputs strongly drive motor cortical activity, what signals do these inputs provide? In order to address this question, we again silenced motor cortical activity in VGAT-ChR2-EYFP mice, putting the motor cortical network - and thus the contribution of recurrent dynamics - in the same initial state at the end of the perturbation on different trials. When we removed the motor cortical inactivation, we allowed the network to recover on some trials, but immediately silenced the thalamus on other trials. This thalamic inactivation blocked reaching following the removal of motor cortical suppression, and the animal frequently reached to the target following the removal of thalamic inactivation (fig. 4d; supplementary fig. 9a). These post-thalamic-inactivation reaches were generated by the same neural pattern that drove reaching on control trials and post-motor-cortical-inactivation trials (supplementary fig. 9b). Thalamic inactivation did not act by merely silencing motor cortical spiking; firing rates during this epoch fluctuated extensively (fig. 4e; supplementary fig. 9c). Population activity with and without thalamic inactivation began in the same initial state, but rapidly diverged after the end of motor cortical suppression (fig. 4f). This indicates that external inputs drove the motor cortical activity when the motor cortex was released and thalamus was not perturbed (fig. 4g). We estimated the difference in external inputs between the two conditions (see methods), revealing dimensions of neural activity with a strong influence of external input signals (fig. 4h). These input differences were not step functions, but were multi-phasic within the first 200 ms following the end of motor cortical inactivation. This suggests that external inputs do not merely provide tonic drive, but rather shape the time-varying pattern of motor cortical activity required to generate reaching.

## Discussion

Using a combination of electrophysiology, optogenetic perturbations, and behavioral tracking, we found that the motor cortex of the mouse, like the primate motor cortex, is a dynamical system that produces a time-varying pattern controlling the arm. We then used this experimental preparation and the dynamical systems framework to dissect, for the first time, the neural mechanism that generates this pattern. We found that the motor cortex is not an autonomous pattern generator, in which strong recurrent dynamics produce the output; rather, it must be driven strongly by external input during the initiation and execution of movement.

How do local dynamics and inputs interact to produce the cortical output pattern? In the extreme case of an autonomous system, the entire pattern can be produced without input (fig. 1b), while at the other extreme, the pattern may be inherited almost entirely from the inputs (fig. 1c). Our results rule out alternatives near the extreme of autonomy: when inputs are removed, the motor cortex does not merely produce a corrupted or scaled-down version of the normal pattern, but instead moves to a new fixed point (fig. 4f). Most likely, the local dynamics and inputs interact to produce this pattern. Furthermore, it is possible that this interaction is nonlinear; that is, the evolution of cortical activity over time might be described by d/dt r(t) = ϕ(r(t),u(t)).

Studies in nonhuman primates have emphasized the role of external inputs in setting up the appropriate initial state in motor cortex for movement generation, and have suggested that motor cortex is largely autonomous during movement execution. By contrast, after we set motor cortex to three distinct aberrant initial states, motor cortex rapidly generates the appropriate pattern for reaching (fig. 2c) without returning to the initial state observed on control trials (supplementary fig. 4). This is consistent with previous results showing that in a mid-movement switching task, cortex can bypass the normal preparatory state (*27*). Inactivation of external inputs to motor cortex prevents progression from aberrant states to the activity pattern driving reaching (fig. 4d-f). Thus, a specific initial state in motor cortex is not required for movement generation; rather, external inputs are capable of compensating for aberrant initial states and driving the appropriate pattern.

What information do external inputs convey? When we inactivated motor cortex in the middle of a reach, we removed trial-to-trial variability in neural activity during the laser-on epoch (supplementary fig. 6d), so that any information about differences in hand position on different trials was erased from motor cortex. Following this perturbation, we decoded hand velocity from neural activity (fig. 2e), and found that pairs of trials with similar observed velocity profiles had similar decoded velocity profiles (fig. 2f). This suggests that upstream brain regions have information about the current and target state of the arm, then route this information through motor cortex to compensate for the perturbation. Some of this information might come from relatively direct proprioceptive relays through the thalamus. An intriguing possibility, however, is that signals about the state of the arm and the target might be maintained through persistent activity in another area, such as the frontal cortex on the same side, or by the contralateral cortex.

Motor cortical inactivation in the absence of a cue often produced post-perturbation neural pattern for movement, along with a coordinated reach to the target. One explanation for this phenomenon might be that the intrinsic motor cortical firing rate derivative, h(r(t)), was large in the inactivated state (due, for example, to hyperpolarization-activated conductances), and was able to initiate the normal pattern for reaching without any external input, whether or not a cue had been presented previously. The observation that silencing thalamus after motor cortical inactivation blocked post-motor-cortical-inactivation reaching, however, suggests that intrinsic motor cortical factors, h(r(t)), are inadequate to generate the appropriate pattern. Instead, perturbation of motor cortex may initiate the command for reaching in other brain areas, and this command may be unmasked when a reach is initiated following the end of the perturbation.

Previous studies of motor cortex have suggested that it operates autonomously during reaching, much like the spinal networks for locomotion: once the pattern has been initiated, strong recurrent dynamics within the local network complete it (*2, 22, 25*). By contrast, we have shown that this pattern can only be produced by strong external input that is maintained throughout the entire movement. In this view, motor cortex integrates signals from diverse brain areas, including the cerebellum and basal ganglia, and translates these signals into the language of the lower motor centers. While motor cortex is a bottleneck for descending motor commands, its activity patterns are crucially molded by its embedding within a vast and distributed network of loops between muscles, the sensory periphery, subcortical regions, and other cortical areas.

## Methods

### Behavioral task and video analysis

Mice were fitted with head posts, food restricted, and trained to reach for food pellets, as described previously (*16*). All data in this manuscript, including those from the behavioral experiments, are previously unpublished. WaveSurfer (http://wavesurfer.janelia.org/) was used to control the behavioral stimuli. Video of the behavior was recorded at 500 Hz using BIAS software (IO Rodeo, available at https://bitbucket.org/iorodeo/bias) and two high-speed cameras (PointGrey, Flea3), which were calibrated to allow 3D triangulation of hand position (Caltech Camera Calibration Toolbox for Matlab, http://www.vision.caltech.edu/bouguetj/calib_doc/). Two types of information were extracted from video: ethograms labeling the frames in which lift, hand open, grab, supination, hand at mouth, and chew occurred, obtained using the Janelia Automatic Animal Behavior Annotator (https://github.com/kristinbranson/JAABA), and the position of the hand in space, obtained using the Animal Part Tracker (https://github.com/kristinbranson/APT). All procedures were approved by the Institutional Animal Care and Use Committee at Janelia Research Campus (protocol 13–99).

### Automatic behavior characterization

Using an adaptation of the Janelia Automatic Animal Behavior Annotator (JAABA) (*28*), we trained automatic behavior classifiers which input information from the video frames and output predictions of the behavior category -- *lift, hand-open, grab, supination, at-mouth*, and *chew*. We adapted JAABA to use Histogram of Oriented Gradients (*29*) and Histogram of Optical Flow (*30*) features derived directly from the video frames, instead of features derived from animal trajectories. The automatic behavior predictions were post-processed as described previously (*16*) to find the first lift-hand-open-grab and supination-at-mouth-chew sequences. For the long-duration thalamic perturbation experiments (fig. 4c, supplementary fig. 8), we used the last lift detected before laser onset for aligning data. Tracking of hand position was performed using the Animal Part Tracker (APT) software package. Hand position was annotated manually for a set of training frames, and the cascaded pose regression algorithm was used to estimate the position of the hand in each remaining video frame.

### Electrophysiological recordings

Neural recordings were performed using the Whisper acquisition system (Janelia Applied Physics and Instrumentation Group) and 64-channel silicon probes (NeuroNexus A4×16-Poly2–5mm-23s-200–177-A64 or Janelia 4×16 probes). These probes consisted of four shanks with 16 contacts at the tip of each, over a depth of 345µm (NeuroNexus) or 320µm (Janelia probes). On the day before the experiment, a small craniotomy was made over motor cortex contralateral to the limb, and a stainless steel reference wire was implanted in visual cortex. During the recording session, the probe tips were positioned at bregma +0.5mm, 1.7mm lateral, and slowly lowered to a depth of ~900µm from the cortical surface, and a silicone elastomer (Kwik-Sil, World Precision Instruments) was applied to seal the craniotomy. At the end of the session, the probe was removed, and the craniotomy was re-sealed with silicone to allow a subsequent session on the following day. Signals were amplified with a gain of 200 and digitized to 16 bits at 25–50 kHz, and spike sorting was performed with JRClust (*31*). Spike sorting code is available at https://github.com/JaneliaSciComp/JRCLUST.

### Optogenetic manipulations

Cell-type specific expression of ChR2 was achieved by either using VGAT-ChR2-EYFP mice expressing ChR2 in inhibitory neurons (Slc32a1-COP4*H134R/EYFP, The Jackson Laboratory), or by crossing a Cre driver line to a Cre-dependent ChR2 reporter mouse, Ai32 (Rosa-CAG-LSL-ChR2(H134R)-EYFP-WPRE, The Jackson Laboratory). Experiments were performed in VGAT-ChR2-EYFP (n = 13), Tg(Tlx3-Cre)PL56Gsat X Ai32 (n = 3), Tg(Sim1-Cre)KJ18Gsat X Ai32 (n = 3), or Tg(Rbp4-Cre)KL100Gsat X Ai32 (n = 2) mice (*32*). Experiments were attempted in three additional mice (VGAT-ChR2-EYFP, n = 2, and Tg(Sim1-Cre)KJ18Gsat X Ai32, n = 1), but were aborted due to the poor quality of the electrophysiological signals. An optical fiber (200 µm or 400 µm, NA 0.39, Thorlabs) was coupled to a 473 nm laser (LuxX 473–80, Omikron Laserage) and positioned 2–4 mm over motor cortex in the head fixation apparatus, as described previously. Five VGAT-ChR2-EFYP mice were implanted with an optical fiber over motor thalamus (bregma −1.1 mm, lateral 1.3 mm, depth 3.3 mm). A blue light emitting diode array was directed at the animal’s eyes throughout the session in order to mask the laser stimulus. Three trial types were used: control trials, in which the cue was presented with no laser stimulation, laser + cue trials, in which both were presented, and laser-only trials, in which the laser was turned on without a cue or food administration. A two-second laser stimulus (40 Hz sine wave) was initiated synchronously with the cue for VGAT-ChR2-EYFP mice, or 200 ms before cue onset for Tlx3-Cre X Ai32 and Sim1-Cre X Ai32 mice. Laser power was calibrated to the minimum level necessary to block reaching in probe experiments in the final days of training; this ranged from 10–50 mW at the fiber tip for VGAT animals, and 0.5–6 mW for Tlx3 and Sim1 animals. In mid-reach interruption experiments, a region of the video frame between the average lift and hand open locations was identified using BIAS software, and a contrast change in this region was used to open the laser shutter for 50–100 ms (for cortical inactivation), or for 2000 ms (for thalamic inactivation). All optogenetic perturbations were unilateral, on the side opposite the reaching arm.

### Data analysis

*Peri-lift firing rates (fig. 1g)*. For each neuron, lift-centered spike trains were smoothed with a Gaussian kernel (σ = 50 ms) and averaged across trials. Lift modulation was assessed using a rank sum test comparing the raw spike counts in a 500 ms window centered at lift +200 ms with counts in a 500 ms window centered at lift −750 ms. Multiple comparisons were corrected using the false discovery rate framework (q < 0.05).

*Trial-averaged principal component analysis (fig. 2c, 3b, E4)*. Peri-lift firing rates were extracted by smoothing the spike trains with a Gaussian kernel (σ = 50 ms for fig. 2c, 3b, E4; σ = 25 ms for fig. 4f), Z-scored using the mean and standard deviation of firing rates on control trials for lift-centered analyses or cortex inactivation trials for laser-centered analyses, and averaged within each trial type for each neuron. The window used was −100 ms to 350 ms around each lift (fig. 2c, E4), −250 ms to +250 ms from the end of cortical inactivation (fig. 3b), −500 ms to +500 ms of the end of the cortical inactivation (fig. 4f, blue), or −500 ms from the start of cortical inactivation to +500 ms from the end of the thalamic inactivation (fig. 4f, green). Principal component coefficients were fit using control trials only for lift-centered analyses (fig. 2c, E4), or cortex inactivation only (fig. 4f), and scores were then extracted for all trial types. For the lift-aligned PCA analyses, a control trial was included if lift and grab occurred within 500 ms following the cue, and a laser trial was included if lift and grab occurred within 500 ms of the end of the laser.

*Gaussian Process Factor Analysis (GPFA) and neural distance (fig. 2d, 3c, E3a, E3d-g, E7a-b, supplementary videos 2, 3)*. Neural population activity was reduced to a five-dimensional latent variable space using GPFA (bin size 20 ms) (*33*). A region of the dataset in which the recordings were stable was selected by finding the time interval and subset of neurons that maximized the quantity (usable neurons) X (usable seconds). The target spatial and neural states were defined using the three-dimensional position of the hand, and the five-dimensional latent variable representation of motor cortical activity obtained using GPFA, respectively. In both cases, the states were sampled at grab times, and the target state was defined to be the central location of the grab-triggered states, computed using convex hull peeling (*34*). Only 50% of the control trials were used to calculate the target states. The euclidean distance from the target was then computed for each trial and time point, and the resulting distance curves were centered either on grab time (fig. E3d-g) or the end of the laser (fig. 2d, 3c, E7a-b). In order to test the robustness of the distance analysis, the distance analysis was also performed using the same procedure with a variable number of GPFA factors ranging from three to seven (fig. E3g), and in the original high-dimensional firing rate space (fig. E3h) using a range of smoothing bandwidths σ = 25, 50, 75, 100, and 125 ms.

*Estimation of difference in external input contributions (fig. 3e, 4h)*. We use the rDSM to calculate the difference in the contribution of external inputs between conditions following the end of cortical inactivation. Suppose we have two types of trial, A and B. In fig. 3e, these types correspond to trials in which a post-laser reach occurred or did not occur, and in fig. 4h, they correspond to trials in which thalamus was perturbed following the end of cortical inactivation and trials in which the thalamus was not perturbed. Let r_A_(t) and r_B_(t) denote the population activity (principal component scores) on these trial types, and suppose the cortical inactivation ends at t = 0.

For t ≤ 0, we have fixed r_A_(t) = r_B_(t) = 0 (see supplementary fig. 6). Let ε > 0. According to the rDSM,

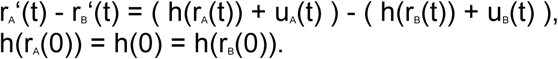

Thus, for small ε,

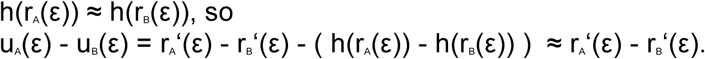

Thus, subtracting the derivative of population activity for the two conditions allows us to estimate the difference in the contributions of external inputs shortly after the end of cortical inactivation.

*Direction of neural trajectories (fig. E4)*. This analysis addresses the question of whether neural activity for post-laser reaches first returned to the control baseline state. For each perturbation (VGAT, Tlx3, and Sim1), we estimated the population activity on control reaches, r_c_(t), and post-laser reaches, r_l_(t), using the first six principal component scores. These captured 98%, 99%, and 97% of the variance in control reaches for VGAT, Tlx3, and Sim1, respectively. At each point in time around the lift, we computed the derivative and divided by the norm of the derivative. This yielded the direction in population activity for control and laser reaches (see fig. E4a), shown by the yellow and blue arrows in fig. (E4b-d). We compared the similarity between the control and laser directions at each point in time by computing the inner product between them. This yielded the yellow curves in the right panels in E4b-d. We then computed the difference between the population activity in the laser condition and the initial control state and normalized it to have length one; this resulted in the direction from the state in the laser condition to the initial control state at each point in time, shown by the red arrows in the left panels of fig E4b-d. We then found the similarity between this direction and the direction of the laser trajectory by computing their inner product, shown in the red curves in the right panels of fig. E4b-d. This analysis suggests that following the perturbations, the neural population recapitulated the pattern for reaching without returning to the control initial state.

*Neural decoding: design of the decoder (fig. 2e, E5)* A linear filter was designed to decode 3D hand velocities from neural activity during reaches. The decoded hand velocities were then used as proxies for the components of neural activity relevant to movement in order to assess the effect of different types of pre-lift perturbations (VGAT, Tlx3, Sim1) on the neural activity during reach (*pre-lift perturbation analysis*), and to assess the variability in the neural trajectories following the post-lift perturbation (VGAT) of motor cortex (*post-interruption variability analysis*).

For decoding, both the 3D hand trajectories (500 Hz, see ‘Behavioral task and video analysis’) and the multiunit neural activity (counts of all detected spikes on each recording channel with 2 ms bins, no single unit identification) were smoothed with the same Gaussian kernel (σ = 25 ms). Velocities were numerically derived from smoothed hand trajectories using a central difference filter of order 8. Firing rates were Z-scored with respect to the activity at rest (computed combining 1.5s windows preceding the start of each trial) and then processed with Principal Component Analysis (PCA) for denoising and dimensionality reduction. Channels with mean absolute Z-scores greater than 100 during movement (e.g. for units with standard deviations very close to zero) were excluded from further analysis.

The decoder uses time-invariant coefficients to decode the hand velocity at any given time as a linear combination of the 15 most recent samples of neural activity (hence up to 28ms in the past) projected to the first few Principal Components. For all datasets analyzed (each corresponding to a different experimental session), PCA was performed on the data matrix obtained combining all control and post-laser neural trajectories (after end of perturbation). The decoder coefficients were obtained by regressing (ordinary least square sense, implemented via QR factorization in Matlab) velocity data against PCA-reduced neural data in a subset of trials (training set). The choice of the trials to include in the training set, and the procedure for choosing the number of Principal Components to use in the decoder, varied slightly between the two types of analysis as described below.

*Neural decoding: pre-lift perturbation analysis* Hand velocities were decoded during reaches (lift-to-grab) which occurred within the first 500ms after the cue (for control trials) or within the first 500ms after the end of perturbation (for laser trials). Within each dataset, the training set used in fitting the decoder coefficients was comprised of a majority of control trials (3/5^th^). An additional subset of control trials (1/5^th^) was used to cross-validate the optimal choice of the number of PCA components to use in the decoder, as follows. For any choice of number of components to keep (up to 95% of variance explained), a decoder was computed from the training control trials and used to decode the hand velocities in the validation control trials. The performance of each decoder was compared in terms of mean squared error (MSE) between observed and decoded velocities. The minimum number of PCA dimensions that guaranteed performance within 1% of the overall minimum MSE across all choices was selected (using this 1% margin guaranteed significant dimensionality reduction in some sessions without compromising decoding performance). Finally, the selected decoder was used to decode hand velocities in the remaining testing subset of control trials (1/5^th^), which were not used for training or cross-validation, and on the laser trials in which reaches occurred at the end of the perturbation.

We found that the decoder performance was not uniform across the three directions (forward, lateral, upward), but was consistently worse in the lateral direction than in the other two directions. This may reflect the smaller extent to which the reaching trajectories sample lateral movements of the hand, or may reflect an insufficient representation in the neural population of motor cortical cells responsible for the lateral movement of the hand (note that lateral movements are more shoulder-dependent than forward and upward movements). We thus showed the decoding performance in each direction separately, and used the R² of the linear regression between observed and decoded hand velocities in each direction as the summary performance metric.

In most of the datasets (all of the VGAT and Tlx3 experiments, and one of the Sim1 experiments), the decoded hand velocities were closer to the observed ones in the control testing trials than in the laser trials (supplementary fig. 5). Since the neural trajectories in laser trials do not exactly follow those in control trials (but are somewhat shifted versions of them, fig. 2c), but the hand trajectories appear qualitatively identical in the two conditions, this suggests that the transformation of motor cortical commands into hand movements was partially modified by the pre-lift perturbation of motor cortex. Subcortical circuits may have played a critical role in compensating for the different levels of cortical output activity in laser trials relative to control. Nevertheless, in most sessions the decoder trained only on control trials still performed reasonably well on laser trials (at least in some of the directions), confirming that the cortical trajectories produced after pre-lift perturbation are qualitatively similar to the normal cortical trajectories (as also observed in the comparison of the PCA trajectories).

#### Neural decoding: post-interruption variability analysis

To investigate the relation between the variability in hand trajectories and that in neural trajectories following post-lift cortical perturbation, we decoded hand velocities from neural activity in the period from 30 ms to 200 ms after the end of such perturbation. The first 28 ms immediately following the cortical perturbation were excluded because the neural data used for decoding would have overlapped with the period of cortical perturbation.

Because the number of post-lift perturbation trials in each dataset was limited (as they usually occurred every 5^th^ control trials), and post-interruption trajectories were qualitatively different from each other and from control trials (because the hand typically drifts to atypical locations during the cortex perturbation), we used a leave-one-out strategy to train a specific decoder for each post-interruption trial. For each of these trials, the decoder coefficients were computed from a training set comprised of all the other post-interruption trials and a large subset of control trials (2 out every 3 control trials, lift-to-grab). The remaining subset of control trials (1/3^rd^) was used to cross-validate the choice of the number of PCA components to use in the decoder, following the same procedure described for the *pre-lift perturbation analysis*.

The post-interruption hand velocities in each trial were then decoded with the trial-specific decoder (which was trained without using any of that trial’s data). As in the *pre-lift perturbation analysis*, the decoder performance was evaluated in terms of the R² of the linear regression between observed and decoded hand velocities in each direction, and was consistently worse in the lateral direction than in the other two directions.

One method to verify if the post-interruption neural trajectories in motor cortex reflect the variability in the hand trajectories is to test whether the similarity (or lack thereof) between the observed hand velocities in any two trials is predictive of the similarity (or lack thereof) in the hand velocities decoded from neural activity in the same trials. For each trial, we computed the 85-dimensional vectors corresponding to the observed and decoded velocities in the window 30–200ms after cortical perturbation (sampled every 2ms), in each of the three dimensions. For each pair of trials, and in each direction, we computed a “dissimilarity score” between the observed hand velocities by taking the L1-norm of the difference between the corresponding 85-dimensional vectors (“Manhattan distance”), and dividing by the number of samples. The resulting score is equivalent to the mean sample-wise distance between the velocity profiles in the two trials. In the same way, dissimilarity scores were computed between the decoded hand velocities in every pair of trials. Finally, we computed the rank correlation (Spearman’s rho coefficient) between the dissimilarity scores in the observed hand velocities and the corresponding dissimilarities scores in the decoded hand velocities (fig.2f). We found a significant positive correlation (p<0.001, one-tailed test) in 6 of the 7 datasets for the directions better decoded (forward and upward) and in 4 of the 7 datasets for the lateral direction.

All analyses were performed with custom-written software in Matlab, except where otherwise noted.

### Code and data availability

Code for automatic annotation of behavior and behavioral waypoint estimation is available at https://github.com/kristinbranson/JAABA. Code for hand tracking is available at https://github.com/kristinbranson/APT. Code for spike sorting is available at https://github.com/JaneliaSciComp/JRCLUST. Data for the present study and code used to analyze data are available from the corresponding author on reasonable request.

## Supplementary video captions

Video 1: Head-fixed prehension behavior and hand tracking. The video shows raw images from two cameras capturing the movement sequence, along with the triangulated three-dimensional location of the hand.

Video 2: Neural population activity and hand position during control and post-laser reaching, centered on cue. Neural state, computed using GPFA, is shown in the left panel, and hand position is shown in the right panel. Each point corresponds to a single trial, with yellow indicating control and blue indicating post-laser reaches. Lift and grab times are green and magenta, respectively.

Video 3: Neural population activity and hand position during control and post-laser reaching, centered on grab. As in video 2, but trajectories are aligned to the grab time.

## Supporting information

## Acknowledgements

B.S., J.G., and A.H. designed the experiments. B.S. and J.G. performed electrophysiological recordings. J.G. performed behavioral experiments. B.S. analyzed electrophysiology and behavior data. J.G. and W.G. analyzed behavior data. M.M. developed the hand velocity decoding analyses. N.V. and K.B. developed preliminary analyses for decoding of behavioral waypoints. M.K. and K.B. developed computer vision algorithms and software. B.S., A.H., M.M. and K.B. wrote the paper with input from all authors. A.H. and K.B. supervised the project.

We thank Byron Yu and the Yu Lab, Steve Edgley, Joshua Dudman, Albert Lee, Brett Mensh, Misha Ahrens, Jeremy Cohen, Arseny Finkelstein, James Fitzgerald, and Kevin Shan for discussions and comments on the manuscript; Allen Lee for tracking software; Tim Harris, Brian Barbarits, Bill Karsh, Steve Sawtelle, Peter Polidoro, and Dan Flickinger for instrumentation development and support; Adam Taylor for development of WaveSurfer; James Jun for spike sorting software; Sammie Chung for assistance with video labeling; Salvatore DiLiso for fiber implantation surgeries; Julia Kuhl for the mouse drawing; and the Janelia vivarium and scientific computing core facilities for support.

---

**Figure S1:**
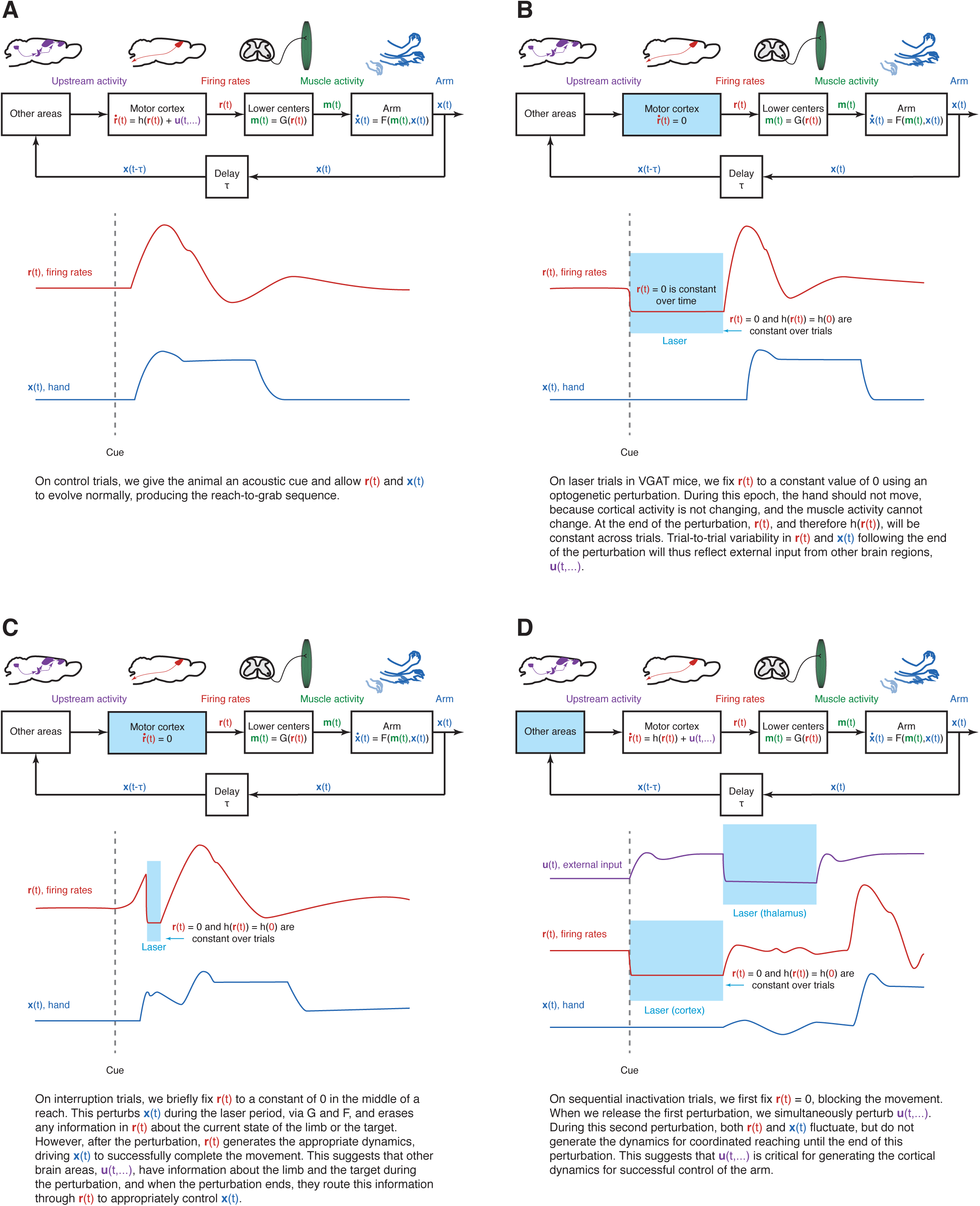
Schematic of experimental design and results.

**Figure S2:**
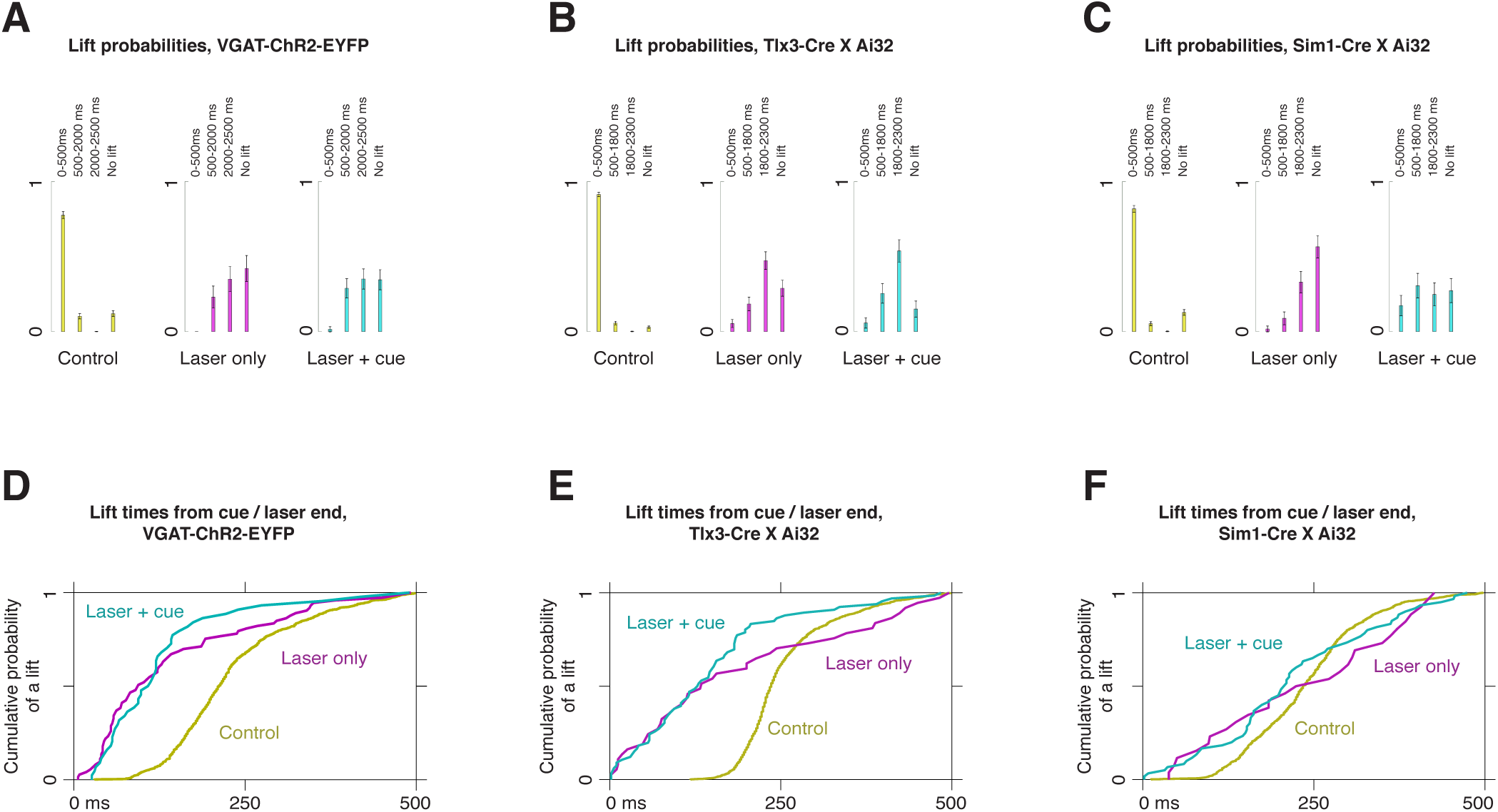
Summary of behavioral effects of optogenetic perturbations. **a**, Probability of a lift in each time bin for control (yellow), laser-only (magenta), and laser + cue (blue) trials for VGAT-ChR2-EYFP mice. The “no lift” bin corresponds to trials on which no lift occurred at any time in the trial. Time 0 corresponds to the start of the trial. **b**, Lift probabilities for Tlx3-Cre X Ai32 mice. **c**, Lift probabilities for Sim1-Cre X Ai32 mice. **d**, Distribution of lift times for trials in which a lift occurred within 500 ms of either the cue (for control trials, yellow), or following the end of the laser (for laser + cue trials, blue, and laser only trials, magenta) in VGAT-ChR2-EYFP mice. On average, the time from the end of the laser to a reach (on perturbation trials) was shorter than the time from the cue to a reach (on control trials). **e**, Distribution of lift times from cue or laser end in Tlx3-Cre X Ai32 mice. **f**, Distribution of lift times from cue or laser end in Sim1-Cre X Ai32 mice.

**Figure S3:**
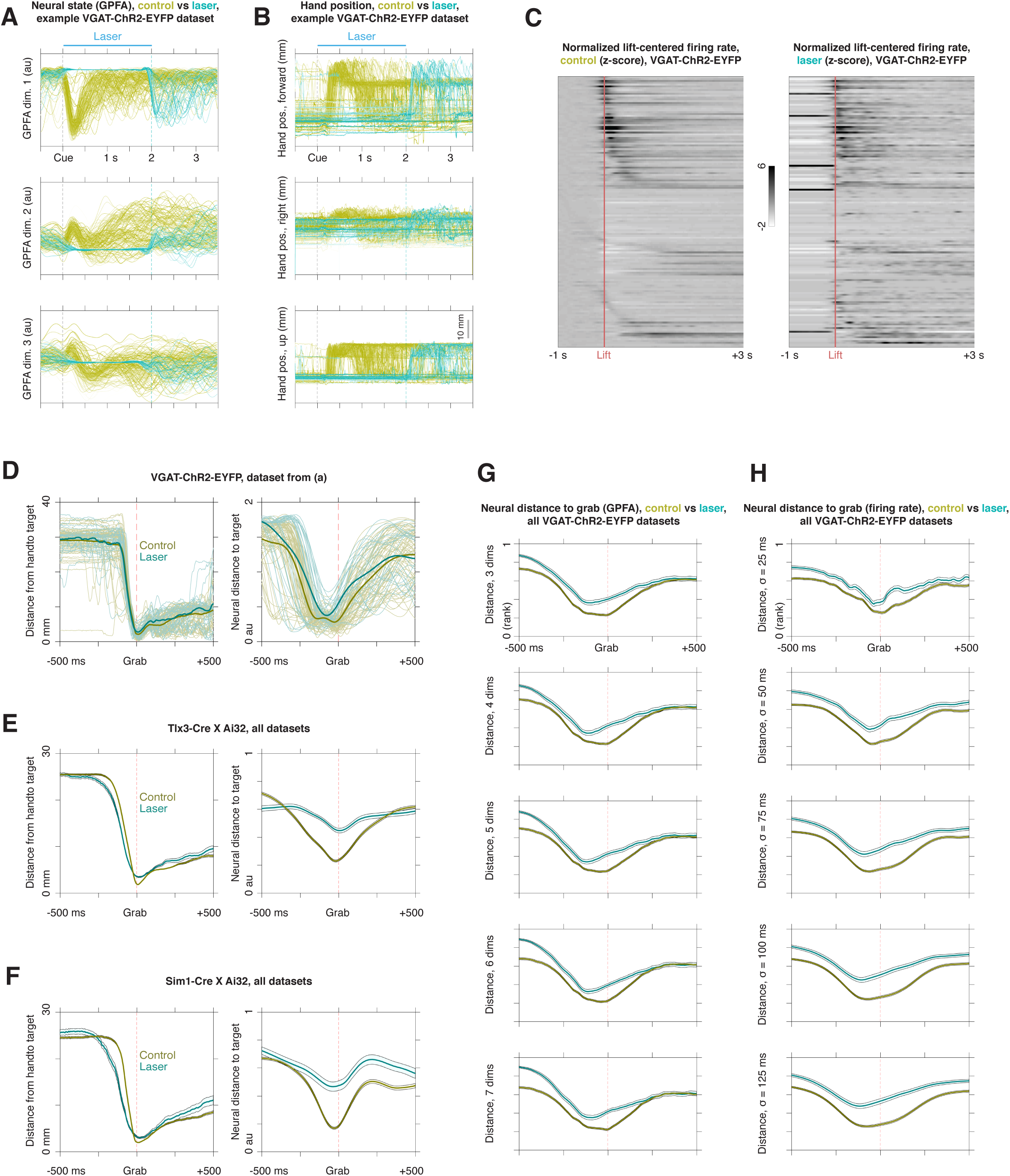
Behavior and neural population activity during reaches on control trials and reaches following the end of motor cortical perturbations. **a**, Single-trial neural activity on control (yellow) and laser (blue) trials estimated using GPFA in an example dataset from a VGAT-ChR2-EYFP mouse. Each panel shows one of the first three GPFA factors. **b**, Single-trial hand trajectories for the same dataset. **c**, Peri-lift firing rate Z-scores for control reaches (left) and post-laser reaches (right) from all neurons recorded in VGAT-ChR2-EYFP mice. **d**, Left: spatial distance from hand to target on control and laser trials, centered on grab, for the dataset in **a** and **b**. Right: neural distance from population state to target state, estimated using Gaussian Process Factor Analysis (GPFA). **e**, Average spatial (left) and neural (right) distance to target on control (yellow) and laser (blue) trials for Tlx3-Cre X Ai32 mice. **f**, Average spatial (left) and neural (right) distance to target on control (yellow) and laser (blue) trials for Sim1-Cre X Ai32 mice. **g**, Average neural distance to target for all VGAT-ChR2-EYFP datasets, obtained using GPFA with a number of dimensions ranging from 3–7; n = 5 mice, n = 7 sessions. **h**, Average neural distance to target for all VGAT-ChR2-EYFP datasets, obtained using firing rate distance over a range of smoothing bandwidths.

**Figure S4:**
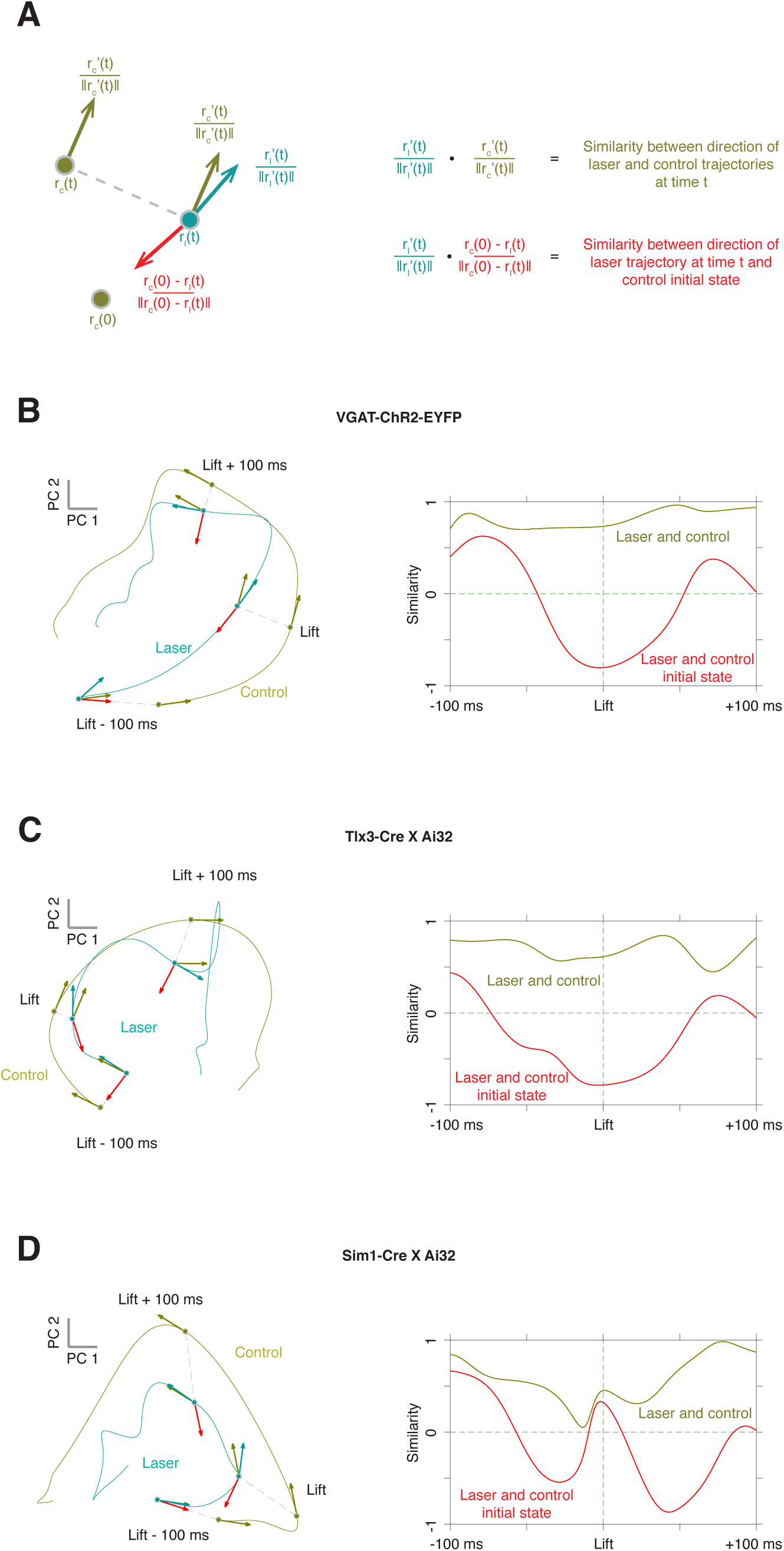
Comparison of the direction of neural trajectories for post-laser reaches with the direction of control trajectories, and with the direction to the initial cortical state on control trials. **a**, Explanation of the analysis method. We represent the population trajectory on control trials, r_c_(t), and laser trials, r_l_(t), using the first six principal component scores, which account for 98%, 99%, and 97% of the variance on control trials for VGAT, Tlx3, and Sim1, respectively. For each time point along the peri-lift neural trajectory r_l_(t) for post-laser reaches, we obtain the direction of the neural trajectory by computing the derivative and dividing by the norm of the derivative (blue). We perform the same calculation for the control trajectory r_c_(t) (yellow), and also compute the direction from the neural state in the laser trajectory to the initial control state (red). We then compare the direction of the laser trajectory with the control direction and the direction to the initial control state by taking the inner product with each. **b**, Left: neural population trajectories (first two principal components) for control (yellow) and post-laser (blue) reaches in VGAT-ChR2-EYFP mice. The direction of the trajectories for control (yellow arrows) and laser (blue arrows) trajectories are shown, along with the direction from the laser trajectory to the control initial state (red arrows). Right: similarity (inner product) between the direction of the laser trajectory and the direction of the control trajectory (yellow curve), and similarity between the direction of the laser trajectory and the control initial state (red curve). **c**, As in **b**, but for Tlx3-Cre X Ai32 mice. **d**, As in **b**, but for Sim1-Cre X Ai32 mice.

**Figure S5:**
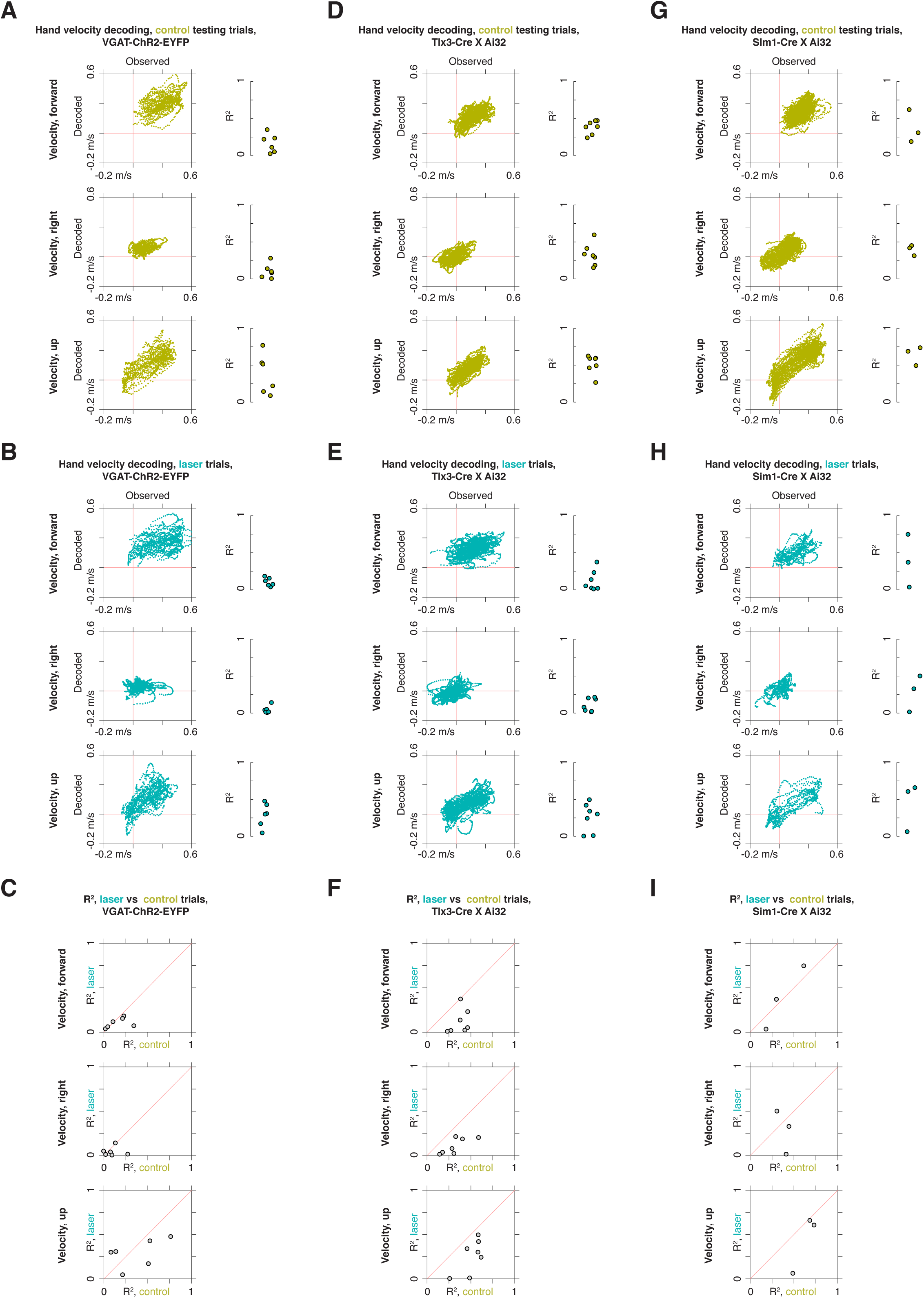
Decoding of hand velocity from motor cortical activity on control and postperturbation reaches. **a**, Left: scatterplots of decoded vs observed hand velocity in the forward, right, and upward directions on control reaches in VGAT-ChR2-EYFP mice in the example dataset from fig. E3**a-b**. Only testing trials not used for training the decoder were used. Right: R2 values for the regression of observed on decoded velocities for control reaches in each VGAT-ChR2-EYFP dataset having at least two post-laser reaches (n = 4 mice; n = 6 sessions). **b**, Left: scatterplots of decoded vs observed hand velocity for post-laser reaches in the dataset from **a**. Right: R2 values for the regression of observed on decoded velocities for post-laser reaches in each VGAT-ChR2-EYFP dataset. **c**, decoding performance for control vs laser. **d-f**, Decoding results for control and post-laser reaches, as in **a-c**, but for experiments in Tlx3-Cre X Ai32 mice (n = 3 mice; n = 7 sessions). **g-i**, Decoding results for control and post-laser reaches, as in **a-c**, but for experiments in Sim1-Cre X Ai32 mice (n = 2 mice; n = 3 sessions).

**Figure S6:**
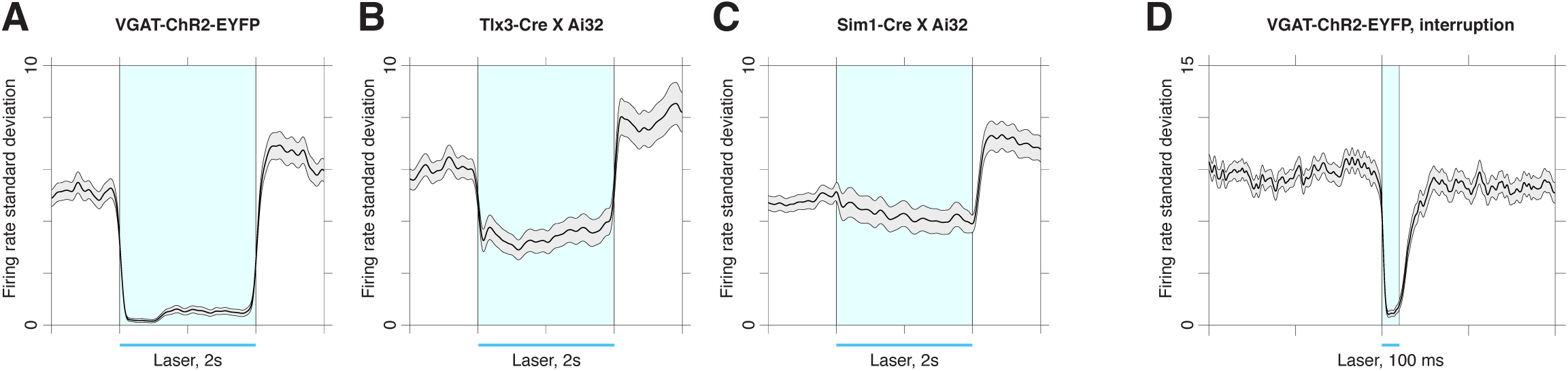
Variability of firing rates during optogenetic perturbations. **a**, Standard deviation of firing rates across trials during laser stimulation in VGAT-ChR2-EYFP mice. The black curve is the standard deviation (over trials), averaged over all neurons. Error bars show standard error of the mean. Identified inhibitory neurons, which exhibited a firing rate increase during the laser, were excluded. Smoothing was applied with a 50 ms Gaussian kernel for each trial. **b**, Standard deviation of firing rates across trials during laser stimulation in Tlx3-Cre X Ai32 mice, as in **a**. Because it wasn’t possible to identify inhibitory neurons, all cells were included in this calculation. **c**, Standard deviation of firing rates across trials during laser stimulation in Sim1-Cre X Ai32 mice, as in **a**. Because it wasn’t possible to identify inhibitory neurons, all cells were included in this calculation. **d**, Standard deviation of firing rates across trials during mid-reach laser stimulation in VGAT-ChR2-EYFP mice, as in **a**. Identified inhibitory neurons were excluded. Because of the short duration of the stimulus (100 ms), the spike trains were smoothed with a 10 ms Gaussian kernel.

**Figure S7:**
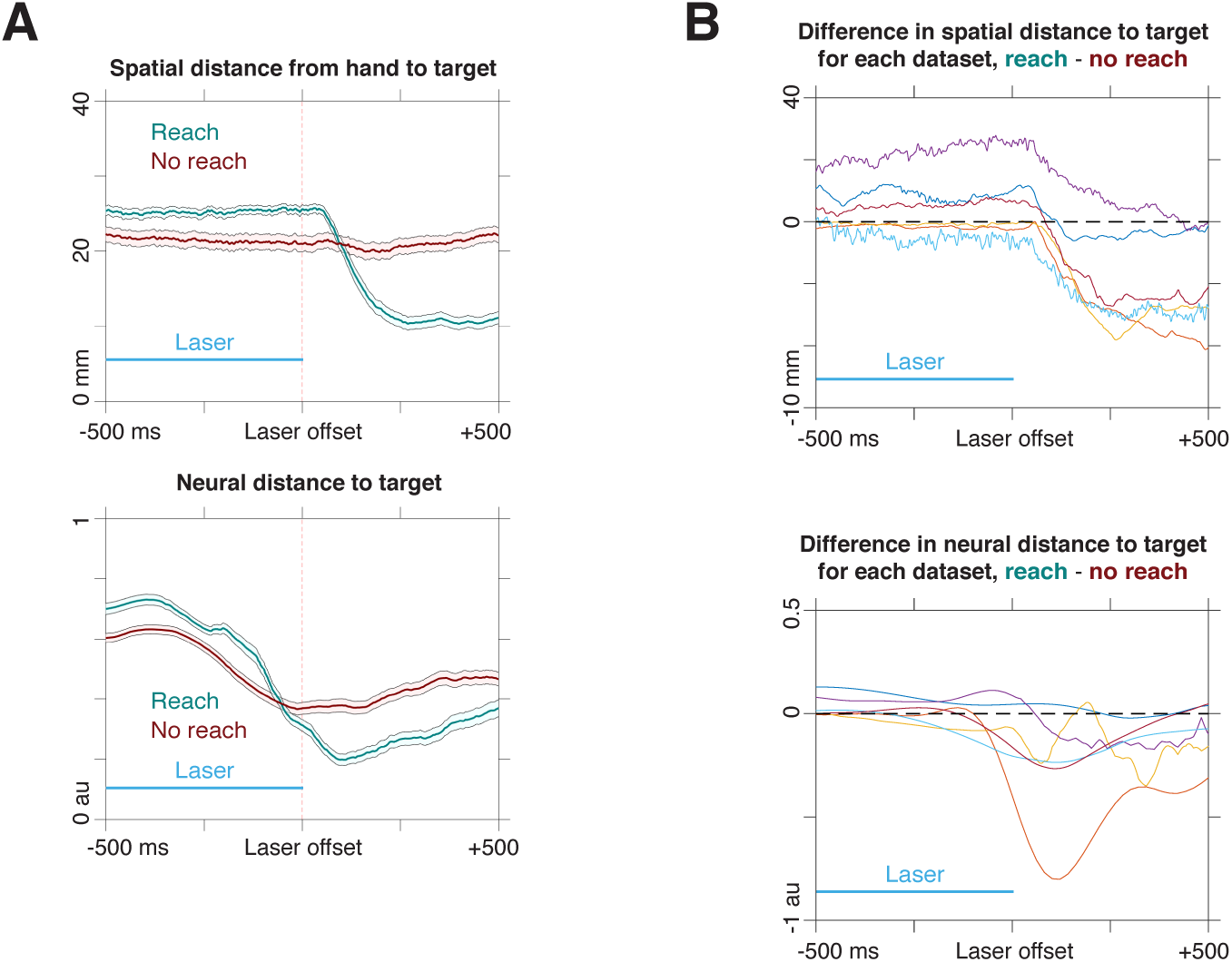
Post-laser distance to grab for trials with and without a post-laser reach in VGAT-ChR2-EYFP mice. **a**, Laser-offset centered spatial (above) and neural (below) distance to target for all datasets; n = 4 mice, n = 6 sessions. One dataset was excluded because no post-laser lifts occurred. The neural state was computed using GPFA with five latent dimensions, and the target state was defined to be the average state at grab on control trials. **b**, Upper: difference between average spatial distance to grab for reach and no reach trials for each dataset (n = 6). Lower: difference between average neural distance to grab for reach and no reach trials for each dataset.

**Figure S8:**
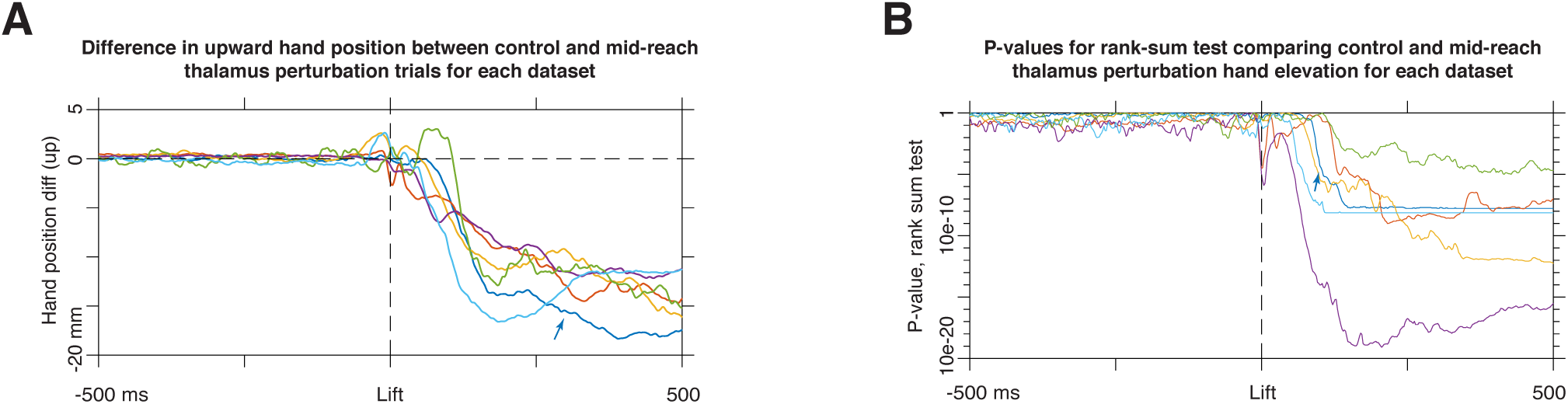
Effect of mid-reach thalamic perturbation on hand trajectory in VGAT-ChR2-EYFP mice. **a**, Difference in hand elevation between mid-reach perturbation trials and control trials for each dataset. The example dataset shown in fig. 4c is marked with the blue arrow. **b**, P-values from rank sum tests at each time point comparing the upward hand position on control and mid-reach thalamic inactivation trials; n = 4 mice, n = 6 sessions.

**Figure S9:**
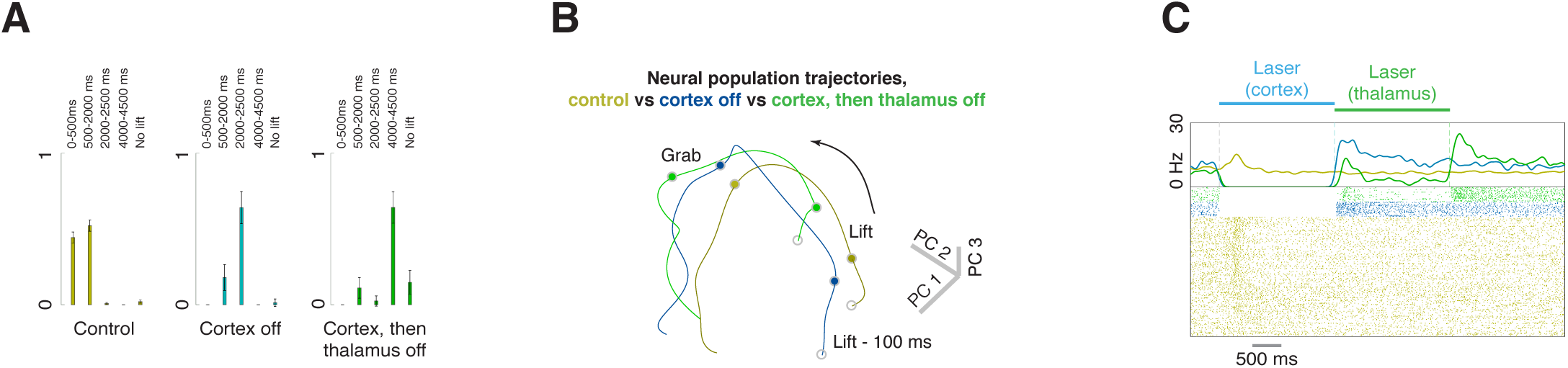
Sequential inactivation of cortex and thalamus. **a**, Fraction of trials with lifts in each epoch for control trials (yellow), cortical inactivation only (blue), and sequential inactivation of cortex and thalamus (green). **b**, Lift-locked neural population activity from lift −100ms to lift +350 ms on control (yellow), inactivation of cortex (blue), and sequential inactivation of cortex and thalamus trials (green), obtained using trial-averaged principal component analysis; n = 3 mice, n = 4 sessions, n = 127 neurons. **c**, Firing rates and spike rasters for an example cortical neuron on control trials (yellow), cortical inactivation (blue), and sequential inactivation of cortex and thalamus (green).

## Notes

#### Summary of Updates

We have updated the manuscript to focus on the question of whether motor cortex is an autonomous or input-driven dynamical system. We have added extensive new experiments and analyses to support the new conclusions.

